# Automatic deep learning-based segmentation and quantification of stented arterial cross-sections for morphometric analysis

**DOI:** 10.64898/2026.04.28.721259

**Authors:** Max Kraftberger, Kristin Spirgath, Tobias Haase, Rubens Bandelin, Tom Meyer, Noah Jaitner, Heiko Tzschätzsch

## Abstract

Arterial vascular diseases, such as atherosclerosis, are among the most serious global health threats. In preclinical studies, morphometric analysis of histological arterial cross-sections is considered the gold standard for assessing vascular remodeling and the effectiveness of therapeutic interventions. However, morphometric analysis is usually performed manually, which is time-consuming, subjective, and requires significant user interaction.

This paper presents a fully automated, operator-independent framework for the precise morphometric analysis of stented arterial cross-sections, extending the previously developed qHisto (quantitative histology) framework for the quantification of various histological components. A neural network for the segmentation of arterial structures was trained and evaluated using 819 cross-sections. In addition, a quantitative analysis of vascular morphology, fibrin area, and lumen asymmetry was performed using 72 cross-sections from coated and uncoated balloons.

The model achieved high segmentation accuracy with a median Dice similarity coefficient of 0.892–0.996. Compared to manual evaluation, the system reduces analysis time by 90%, enabling efficient processing of large datasets. Furthermore, morphometric analysis with qHisto showed significant differences between coated and uncoated balloons, e.g. regarding lumen area (AUC = 0.86) and fibrin ratio (AUC = 0.94).

Our developed framework enables fully automated, comprehensive and standardized analysis of histological arterial cross-sections. This helps to reduce time-consuming, repetitive manual assessments and thus facilitates research of disease mechanisms and treatment effects in preclinical studies.

## 1. Introduction

Cardiovascular diseases, including atherosclerosis, remain a leading cause of mortality worldwide [1-5]. Progressive plaque accumulation within the arterial lumen can reduce or obstruct blood flow and impair cardiovascular function [4, 6]. Endovascular interventions such as stent implantation and balloon angioplasty are established first-line treatments for restoring perfusion in stenotic arteries [5]. However, mechanical vascular injury associated with these procedures provoke an excessive wound healing response, resulting in in-stent restenosis [5, 7, 8]. In arteries affected by in-stent restenosis, neointima formation leads to a significant thickening of the intima [9]. The intima is the inner part of the arterial wall and encloses the vascular lumen [4, 10]. It is separated from the media by the internal elastic lamina (IEL), while the media is separated from the adventitia by the external elastic lamina (EEL) [4, 10]. To mitigate restenosis, stents and balloons are coated with anti-proliferative or anti-inflammatory drugs, such as paclitaxel or sirolimus [7, 8].For the investigation of vascular remodeling processes and the development of endovascular therapeutic strategies such as drug-eluting stents and drug-coated balloons, preclinical studies commonly employ in-stent stenosis models in species, including mice [11], rats [12], rabbits [13], and pigs [14]. Particularly porcine models are widely used for the translation of novel endovascular therapies to clinical application [15], due to their close anatomical and physiological similarity to humans [16]. Beyond *in vivo* imaging modalities, histomorphometric analysis of stained arterial cross-sections is considered the gold standard for detailed assessment of vascular remodeling and treatment efficacy. Morphological parameters typically assessed are measurements of luminal, neointimal, and medial areas, cross-sectional lengths, and soring of fibrin [8, 17, 18].

Conventional morphometric analysis is usually performed manually by delineating anatomical structures, making it highly time-consuming and dependent on operator expertise. Even experienced investigators require 10–15 minutes to fully annotate a single cross-section, limiting scalability for large datasets. Moreover, manual analysis is prone to inter-operator variability arising from subjective interpretation, fatigue, and heterogeneity in tissue quality, staining, and anatomical features [19, 20].

Computer-aided analysis using algorithmic methods can potentially increase objectivity and enable a rapid and cost-effective analysis of large amounts of data by enabling operator-independent annotation of medical data [19, 20]. Approaches from previous studies utilize classical image processing and machine learning (ML) methods, such as region-based segmentation approaches or color-based clustering methods with semi-automated contour tracing [21, 22]. However, these approaches typically require user-defined initialization or manual refinement and can therefore be sensitive to staining variability, irregular tissue boundaries, and artifacts [21, 22]. In contrast, deep learning (DL) approaches can learn hierarchical and contextual image features directly from the data and have consistently outperformed traditional image processing and conventional ML methods in challenging microscopy and histology segmentation tasks [23-25]. Recent research by Danilov *et al*. in 2024 showed that DL methods can be used for the segmentation of microvascular structures to analyze vascular tissue regeneration [26]. In this work, we used nnU-Net, a self-configuring segmentation framework that automatically adapts hyperparameters such as pre-processing, architecture, training, and post-processing to a given dataset and has demonstrated state-of-the-art performance across a broad range of biomedical segmentation benchmarks [27]. To the best of our knowledge, no DL approach has yet been developed for segmenting structures in stented arterial cross-sections.

The limitations of traditional histomorphometric analysis are not only related to the manual evaluation but also to the parameters that are currently used for the assessment. Besides measurements of areas and lengths, fibrin deposits are analyzed to assess inflammation and delayed tissue healing. Since manual segmentation of small and discontinuous deposits would be difficult and highly time-consuming, fibrin deposition is traditionally assessed using a semiquantitative scoring system. However, this system is subjective and only considers fibrin deposition around the stent struts and not within the entire neointima and is therefore limited to stented histological cross-sections. As a further parameter, the progression of in-stent stenosis can be characterized by the degree of symmetric or asymmetric neointima formation [28]. However, there is no established parameter yet to evaluate lumen asymmetry. Previously proposed parameters, such as the ratio of minimum to maximum neointima thickness, may yield misleading high values in cases of overall thin neointima thickness [28]. Therefore, the lumen asymmetry in this study is derived from the concept of vessel eccentricity, which is based on centroid displacement [22], and is thus independent of the overall neointima thickness.

For these reasons, we (I) developed a fully automated DL-based pipeline comprising pre-processing, segmentation, and post-processing for comprehensive quantification of established morphological key parameters in arterial cross sections. Additionally, we (II) introduce the fibrin ratio and lumen asymmetry as improved parameters to enhance standardization and robustness. The fibrin ratio, calculated as the ratio between area of fibrin and neointimal area, enables an objective, comprehensive, and quantitative assessment of fibrin deposition which is not limited to stented cross-sections. Furthermore, we validated the pipeline using exemplary data from vessels treated with coated and uncoated balloons. The system is part of the qHisto (quantitative histology) framework, which was developed for the automated segmentation and quantification of various histological components. It is integrated into the BIOQIC-Apps platform of Charité – Universitätsmedizin Berlin, an open-access environment for image processing software, ensuring compatibility and high-performance computing (HPC) cluster access (https://bioqic-apps.charite.de) [29].

## 2. Methods

### 2.1. Image dataset and acquisition

This retrospective study was conducted exclusively using previously acquired digital histological images of arterial cross-sections. No animals were used, handled, or sacrificed specifically for the purpose of this study. The underlying animal experiments from which the tissue samples originated were approved in accordance with the guidelines of the European commission directive 86/609/EEC and the German Animal Protection Act based upon the Animal Ethics Committee approvals (Saxony–Anhalt, Germany). The dataset comprised digital images of stented arterial cross-sections obtained from domestic pigs that underwent coronary or peripheral stent implantation. Arterial segments were embedded in methyl methacrylate, sectioned at 5–8 µm thickness using either microtome or laser cutting techniques, and stained with Movat’s Pentachrome or Masson–Goldner’s Trichrome, respectively. A total of 819 annotated images (arterial wall dataset) were included for the training and evaluation of the arterial wall model, consisting of 666 microtome-cut sections and 153 laser-cut sections. Since strut void annotations were only available for part of this dataset, a subset of 205 images was used for the strut void prediction model (strut void dataset). Additionally, 72 independent images (Coating Dataset) were used to evaluate the model in a practical application scenario and to analyze which parameters enable a reliable differentiation between two treatment groups. The coating dataset comprises 36 images of vessels treated with drug-coated balloons (paclitaxel) and 36 images of vessels treated with uncoated balloons. An overview of the respective datasets and their usage is given in Table 1.

**Table 1.** Overview of datasets used. The table shows the content and use cases of the different datasets used in this study.

| Dataset | Content | Use case |
| --- | --- | --- |
| Arterial wall | 819 scans (655 training, 185 test) | Training of arterial wall model |
| Strut void (subset of arterial wall dataset) | 205 (185 training, 20 test) | Training of strut void model |
| Coating | 72 (36 coated, 36 uncoated) | Validation of diagnostic capability |

Image acquisition was performed using a Hamamatsu NanoZoomer slide scanner at a spatial resolution of 0.228 µm/pixel (111,403 dpi). Image dimensions were approximately 5,500 µm × 5,500 µm for smaller sections and 45,000 µm × 7,000 µm for larger sections. **Figure 5Figure 5Figure 5Figure 5Figure 5Figure 5**

### Ground-truth morphometric assessment

Annotations were generated using NDP.view2 software (Hamamatsu) and included measurements of areas and diameters delineated by the vessel lumen, IEL, and EEL. Manual morphometric analysis performed by one expert observer (TH) was available for the arterial wall dataset and served as the reference standard for model training and evaluation. The following morphometric parameters were assessed in all available data: luminal area, area within the IEL, area within the EEL, lumen diameter, IEL diameter, EEL diameter, maximum neointimal thickness (**Figure 1**). Diameters of the lumen, IEL and EEL were determined based on the delineated contours. For each structure, the longest possible diameter was first drawn across the segmented area. A second line was then placed perpendicular to the first line through its midpoint (**Figure 1**). The mean of these two orthogonal measurements was calculated and recorded as the respective diameter. Maximum neointimal thickness was measured at the location of greatest distance between luminal border and the IEL. Derived parameters included neointimal area (IEL area minus luminal area), medial area (EEL area minus IEL area), neointimal thickness (IEL diameter minus luminal diameter), medial thickness (EEL diameter minus IEL diameter), and lumen loss (neointima area divided by the IEL area).

**Figure 1.**
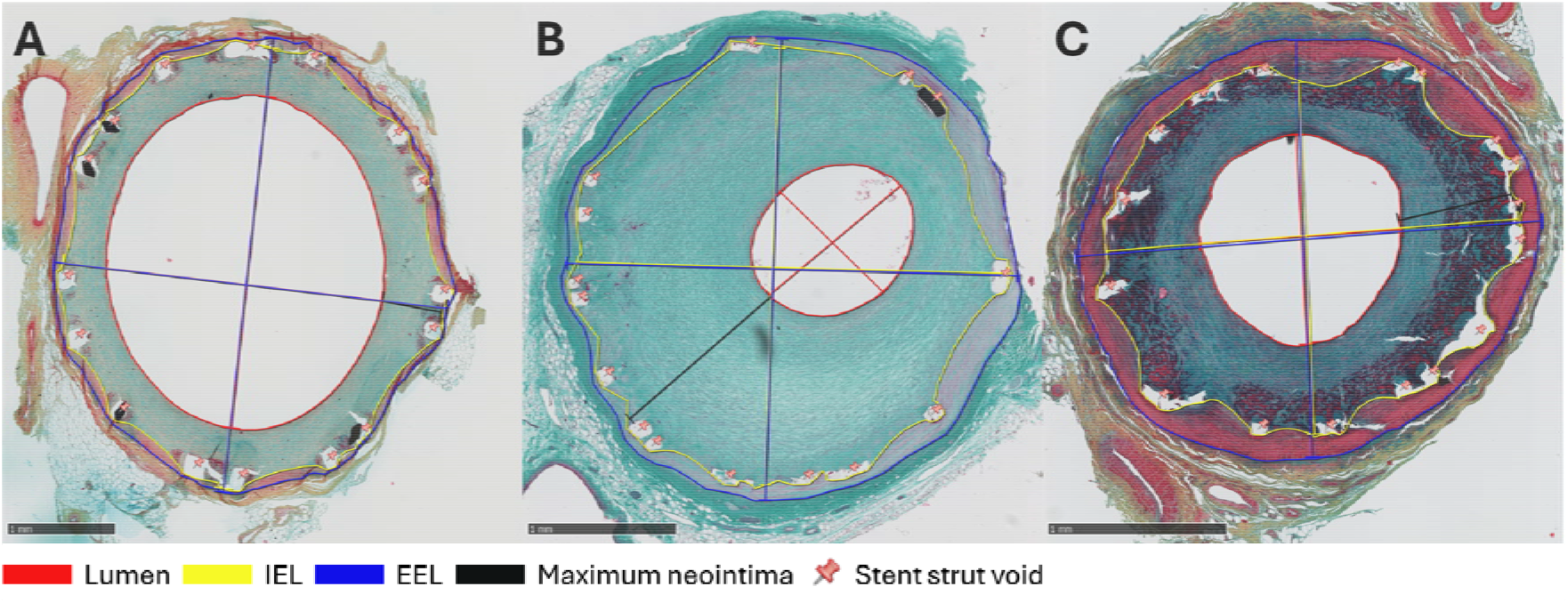
Manual histomorphometric analysis of stented arterial cross-sections representing the ground-truth. Representative Movat’s Pentachrome– (A, C) or Masson–Goldner–stained sections (B) illustrating manual delineation of the area and diameters of the lumen, internal elastic lamina (IEL), and external elastic lamina (EEL). Maximum neointimal thickness was measured at the site of greatest distance between the luminal border and the IEL. Bar = 1mm

In addition to quantitative morphometry, stent strut–associated circumferential fibrin deposition was assessed in the coating dataset using a semiquantitative scoring system: (0) no fibrin; (1) <25% fibrin coverage; (2) 25–50% fibrin coverage; (3) 50–75% fibrin coverage; and (4) 75–100% fibrin coverage. Scores were averaged for all struts within a cross-section to obtain a mean fibrin score per section [8]. Semiquantitative fibrin scores were used to assess the correlation with the automated qHisto fibrin ratio.

The following parameters were considered for diagnostic validation: lumen area, neointimal area, lumen loss, lumen diameter, neointimal thickness, medial thickness, maximum neointimal thickness and fibrin score.

To assess inter-observer variability, 60 out of the 819 images from the arterial wall dataset, consisting of 30 microtome-cut and 30 later-cut cross-sections, were independently morphometrically analyzed by a second observer (MK) using the same protocol.

### 2.3. qHisto

The qHisto framework was originally developed for the automated segmentation and quantification of various histological components of the liver [30]. In this work, qHisto is further extended to enable the segmentation of arterial structures.

The qHisto-framework is integrated into the BIOQIC-Apps website and available for Charité internal use; it can also be used externally without a GPU. This allows the developed qHisto artery pipeline to be executed sequentially upon user request. The fully automatic pipeline begins by converting the uploaded NDPI files to PNG files, followed by an eightfold downsampling to approximately 1.824 µm/pixel in order to minimize memory usage while preserving sufficient structural detail. The images are then used sequentially by the trained models to predict arterial structures and strut voids. Post-processing is then performed to refine the segmentation. Based on the final predictions, areas, diameters, lumen loss, thicknesses, lumen asymmetry, and fibrin ratio are determined. Finally, the predicted structures and the determined diameters are converted to the NDPA file format and saved for convenient human verification. A schematic overview of this workflow is illustrated in Figure 2.

**Figure 2.**
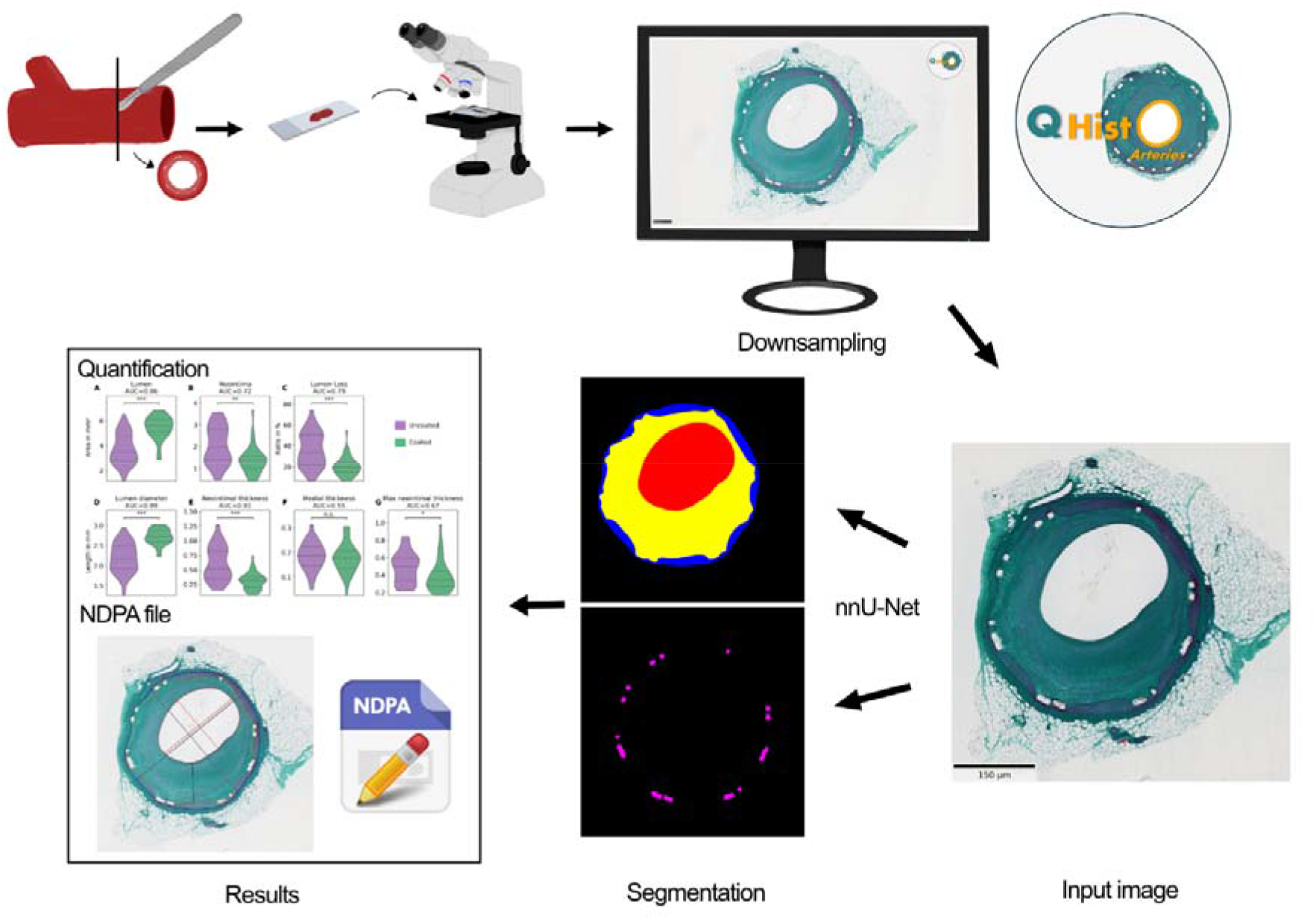
Workflow of the qHisto artery framework. An eightfold downsampling is applied to the high-resolution scan of a stented arterial cross-section. Afterwards the trained models are applied to the image and post-processing steps are performed. Based on the predictions of the arterial wall structures and strut voids the quantification of the areas, diameters, lumen loss, thicknesses, lumen asymmetry and fibrin are conducted. The predictions are additionally saved as an NDPA file.

The DL pipeline was implemented in Python and executed in a virtual environment on a HPC cluster by Charité and BIH (Berlin Institute of Health). The source code of the qHisto framework and the trained model are available in the following GitLab repository: https://gitlab.com/Charite-IMI/qhisto_arteries.

#### 2.3.1. Automatic segmentation

##### 2.3.1.1. Arterial structures

The arterial wall dataset containing expert annotations of the lumen, IEL and EEL. The dataset was randomly divided into 80% training (655 images) and 20% test (164 images) data.

The test dataset was used to train an nnU-Net v2 [27] model for the prediction of the arterial structures. The nnU-net is a self-configuring framework based on the standard U-Net encoder-decoder structure [27]. It automatically adapts hyperparameters such as preprocessing, network architecture, training, and post-processing to the respective dataset [27]. Moreover, five-fold cross-validation was employed, in which the dataset was divided into five parts [27]. Each part served as a validation set once, while the remaining parts were used as training data [27]. Based on the cross-validation results, nnU-Net automatically selected the best performing ensemble of configurations for inference [27]. Training was performed using the 2D U-Net configuration, with all parameters set according to the nnU-Net default settings.

To ensure accurate and high-quality segmentations, certain post-processing steps were applied. These included removing incorrectly identified structures by deleting small objects (empirically determined area of less than 500 pixel (1662 µm^2^)) and objects without adjacent polygons of the correct class. Gaps within the segmentation were closed. Furthermore, it was ensured that the lumen was completely enclosed by the neointima and the neointima by the media. Otherwise, a one-pixel-wide line of the respective label was added. Finally, all objects, except the lumen, that were enclosed by only one class were assigned to that class.

##### 2.3.1.2. Strut voids

The prediction of the strut voids was done by a separate nnU-Net model, since strut void annotations were only available for the strut void dataset, which is a subset of the arterial wall dataset. Due to the much smaller dataset, 90% (185 images) were used as training data, while 10% (20 images) were used as test data. Training was performed using the 2D U-Net configuration with all parameters set according to the default settings of nnU-Net.

Since strut voids can only occur within the arterial wall, a post-processing step was applied. Here, the structures previously predicted by the arterial wall model were used to remove all incorrectly predicted strut voids outside the arterial wall.

##### 2.3.1.3. Fibrin

In order to detect and calculate the fibrin area within the neointima, color intensity values for fibrin regions must be extracted. For this purpose, the color deconvolution plugin from ImageJ was used to extract the color space matrices of the fibrin area for the different stainings. These matrices were then integrated into a custom Python script for color deconvolution. Based on the staining method, the corresponding color space matrix was selected and the processing for the respective staining was performed. The color deconvolution for the fibrin stain was then applied to the masked neointima area to separate the fibrin area. The image was converted to greyscale, and a Gaussian filter (σ = 2 pixel) was applied. Afterwards, an adaptive threshold for binary segmentation of the fibrin regions was used. Additionally, the strut void segmentation mask was applied to remove pixels belonging to strut voids. The fibrin ratio was then calculated by dividing the fibrin area by the neointima area.

#### 2.3.2. Morphological quantification

The areas and diameters for all arterial structures were determined based on the post-processed segmentation maps. The diameters of the respective structures were measured in two perpendicular directions and designated as diameter *a* and *b*. Diameter *a* was implemented as the longest possible distance within the structure. Diameter *b* was implemented as the longest distance within the structure, which is perpendicular to diameter *a* and passing through its center. All derived parameters (neointima area, lumen diameter, lumen loss, neointimal thickness and medial thickness) were calculated as described for conHisto. To calculate the maximum neointima thickness, the nearest lumen point is determined for each IEL point, considering only connections that lie entirely within the neointima. The maximum value is then obtained from these distances. In addition to the used parameters described above, the lumen asymmetry and fibrin ratio were used for diagnostic validation.

#### 2.3.3. Lumen asymmetry

The lumen asymmetry provides information about the concentricity of neointima formation. It was defined as the distance *d* of the lumen centroid to the IEL centroid, divided by the mean radius *c* of the IEL. A higher lumen asymmetry corresponded to a more asymmetric neointima formation.

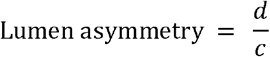

### 2.4. Statistical analysis

To evaluate the model predictions as well as the inter-operator variability for the arterial structures, the Dice similarity coefficient (DSC) was calculated for each test image of the arterial wall dataset. Additionally, the median as well as the 25^th^ and 75^th^ percentiles were computed. The DSC was calculated using the following equation with the segmentation mask of the model prediction *A* and the segmentation mask of the ground truth *B* :

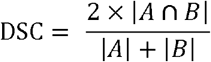

To evaluate strut void segmentation based on the test images of the strut void dataset, each prediction was considered either true positive (TP), false positive (FP) or false negative (FN). If a predicted strut void overlaps with at least one pixel of a ground truth (GT) strut void, it was considered a TP. If a predicted strut void did not overlap with any GT strut void, the prediction was considered a FP. Lastly, if a GT strut void did not overlap with a predicted strut void, it was considered a FN. With these definitions the F1 score was calculated using the formula:

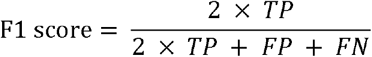

To evaluate the ability of the parameters to differentiate cross-sections from vessels treated with coated or uncoated balloons at stent implantation, the AUC and an unpaired Wilcoxon rank sum test were calculated based on the coating dataset, with *p* values < 0.05 considered statistically significant. To assess the correlation between the qHisto fibrin ratio and the fibrin score from conventional histology (conHisto), the Spearman correlation coefficient was calculated.

## 3. Results

### 3.1. Validation of segmentation models and parameter determination

To evaluate the prediction quality of the trained model, the segmentation results on the 164 test images for arterial structure segmentation were compared with the GT after post-processing and assessed using the DSC. The model achieved high median DSCs for lumen 0.996, [0.991, 0.997], neointima 0.949, [0.896, 0.970], and media 0.892, [0.850, 0.931]. In comparison, a similar median DSCs could be achieved by the inter-operator variability for the lumen 0.997, [0.995, 0.998], neointima 0.986, [0.974, 0.989], and media 0.905, [0.839, 0.928]. **Figure 3** shows representative examples of the best, an average and the worst prediction compared to the GT. In the good and average example, the model achieved a consistent high agreement across all three structures. In the worst case the deviation occurred primarily in the lumen and neointima.

**Figure 3.**
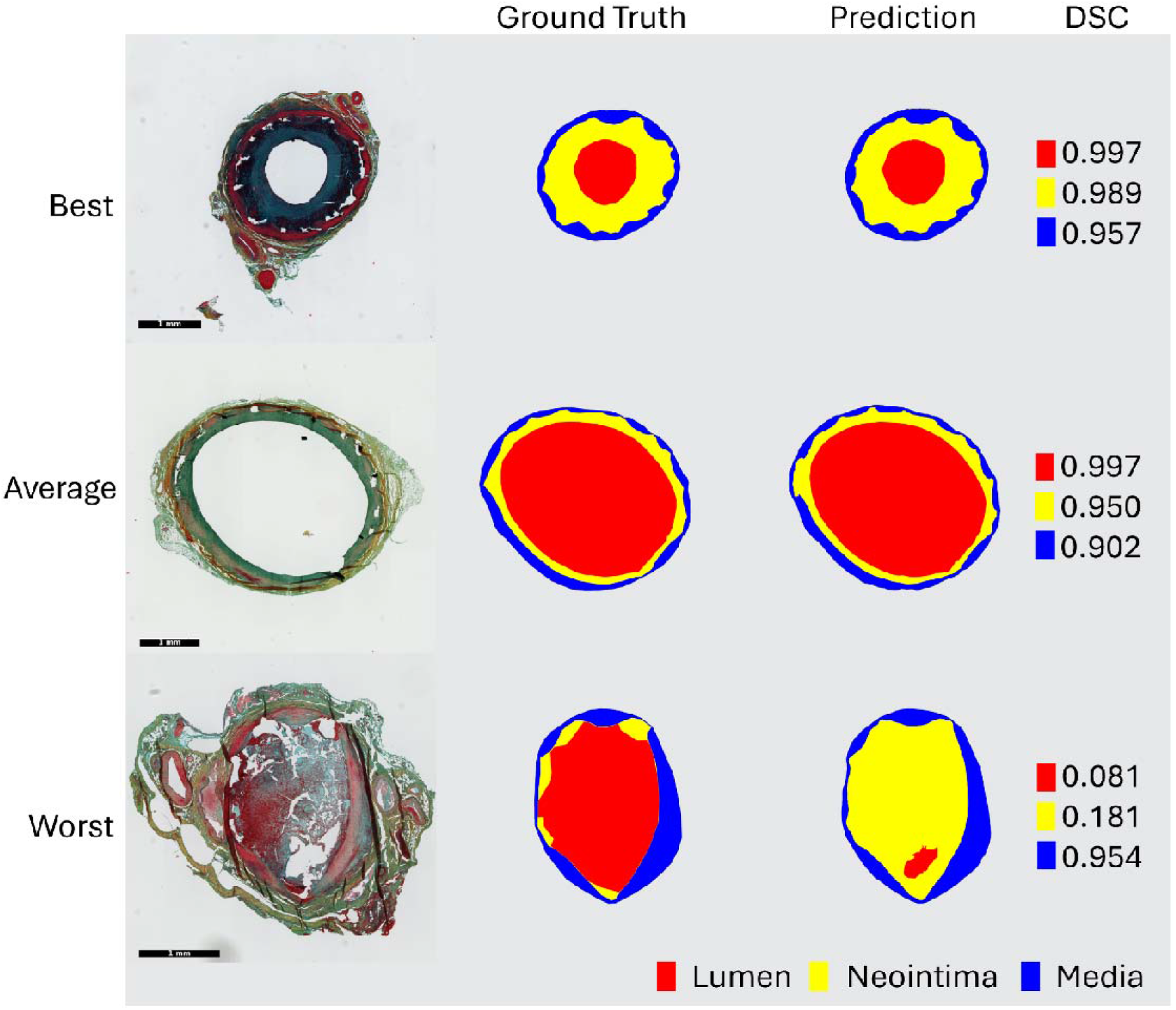
Examples of artery wall structure prediction. Examples of good, average and worst prediction compared to the ground truth. The lumen is shown in red, the neointima in yellow, and the media in blue. The Dice similarity coefficient (DSC) for all structures is shown in the right column. Bar = 1mm.

To evaluate the performance of the strut void model, the segmentation results on the 20 test images for strut void segmentation were assessed using the F1 score. Through this, a median F1 score of 0.97 (IQR: 0.91 – 1.00) was achieved. Representative examples of the best, an average and the worst prediction compared to the GT are shown in **Figure 4**. In the good and average case, a very high F1 score could be achieved, while in the worst case it would be lower.

**Figure 4.**
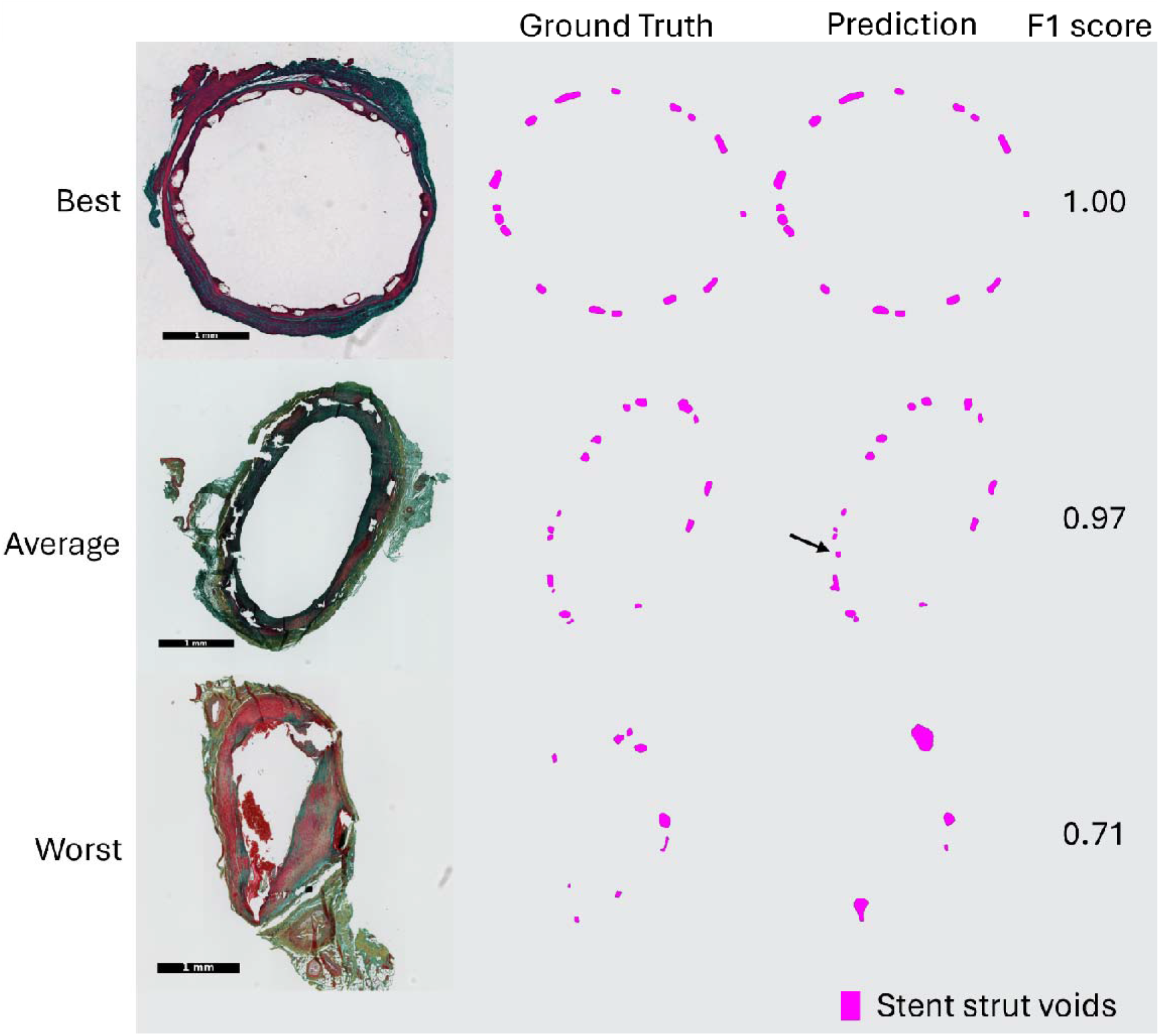
Examples of strut void prediction. Examples of good, average and worst prediction compared to the ground truth. Strut voids are shown in magenta and the respective F1 score is shown in the right column. The arrow indicates false positive prediction. Bar = 1mm.

To evaluate the accuracy of the determined parameters, the relative deviation from the GT was calculated. This yielded in low median relative deviations for the area of the lumen 0% [- 2.48%, 0.34%], neointima -0.05% [-0.67%, 0.93%], and media 0.05% [-0.55% - 0.59%]. Similarly, the median relative deviations are low for luminal, IEL, and EEL diameters and moderate for the maximum neointima thickness (**Table 2: Relative deviations of cross-sectional lengths**. The table shows the median relative deviations of the determined cross-sectional lengths from the ground truth with 25th and 75th percentile in brackets.).

**Table 2:** Relative deviations of cross-sectional lengths. The table shows the median relative deviations of the determined cross-sectional lengths from the ground truth with 25th and 75th percentile in brackets. a: longest distance, b: distance perpendicular to a, IEL: internal elastic lamina, EEL: external elastic lamina.

| Parameter | Relative deviation [%] |
| --- | --- |
| Lumen | 1.24 [0.05, 2.50] |
| Lumen | 0.3 [-0.8, 1.5] |
| IEL | 3.9 [1.8, 7.8] |
| IEL | -0.13 [-2.38, 3.38] |
| EEL | 1.7 [0.8, 3.4] |
| EEL | 0.02 [-1.27, 1.35] |
| Maximum neointimal thickness | 25 [10, 66] |

### 3.2. Demonstration of diagnostic capability

The automated analysis using qHisto distinguished between in-stent segments treated with uncoated and coated balloons (n=36 cross-sections per group) based on multiple morphometric parameters (**Table 3**). The determined classical morphometric parameters are shown in **Error! Reference source not found**.. Strong separation was observed for lumen area (AUC = 0.86, **Error! Reference source not found**.A), lumen diameter (AUC = 0.89, **Error! Reference source not found**.D), and neointimal thickness (AUC = 0.81, **Error! Reference source not found**.E), consistent with well-known drug-related effects of coated balloon treatment on neointimal proliferation [16]. Moderate separation was observed for neointimal area (AUC = 0.72, **Figure 5B**), lumen loss (AUC = 0.79, **Figure 5C**), and maximum neointimal thickness (AUC = 0.69, **Figure 5G**), while medial thickness showed no relevant separation (AUC = 0.55, **Figure 5F**), as expected.

**Table 3.**
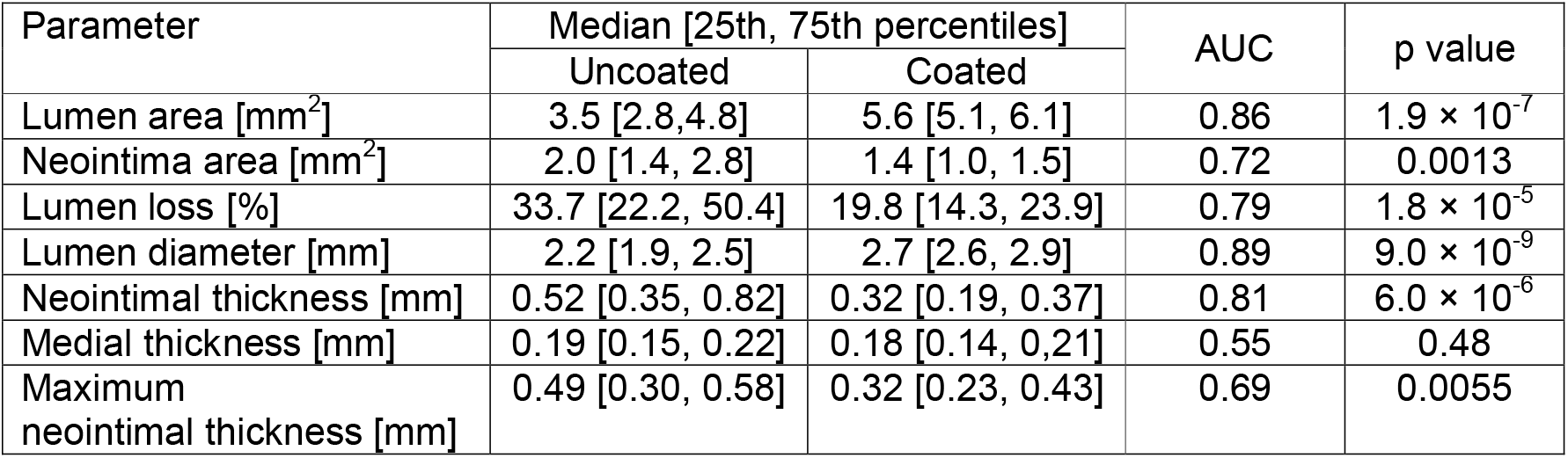
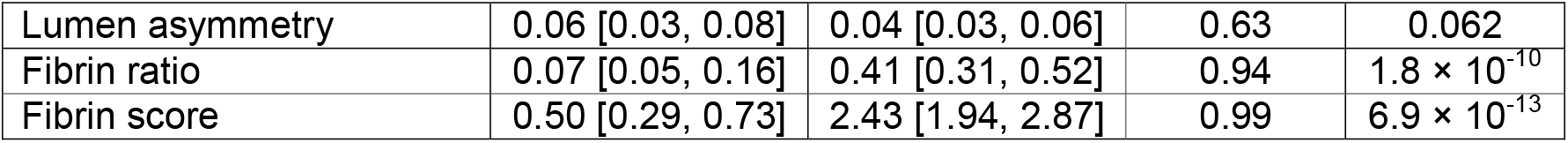
Comparison of parameters between in-stent segments treated with uncoated and coated balloons (n=36 cross-sections per group). The table shows the median values of the parameters with the 25th and 75th percentile in brackets, the AUC (area under the curve), and the p-values.

| Parameter | Median [25th, 75th percentiles] |  | AUC | p value |
| --- | --- | --- | --- | --- |
|  | Uncoated | Coated |  |  |
| Lumen area [mm<sup>2</sup>] | 3.5 [2.8, 4.8] | 5.6 [5.1, 6.1] | 0.86 | $1.9 \times 10^{-7}$ |
| Neointima area [mm<sup>2</sup>] | 2.0 [1.4, 2.8] | 1.4 [1.0, 1.5] | 0.72 | 0.0013 |
| Lumen loss [%] | 33.7 [22.2, 50.4] | 19.8 [14.3, 23.9] | 0.79 | $1.8 \times 10^{-5}$ |
| Lumen diameter [mm] | 2.2 [1.9, 2.5] | 2.7 [2.6, 2.9] | 0.89 | $9.0 \times 10^{-9}$ |
| Neointimal thickness [mm] | 0.52 [0.35, 0.82] | 0.32 [0.19, 0.37] | 0.81 | $6.0 \times 10^{-6}$ |
| Medial thickness [mm] | 0.19 [0.15, 0.22] | 0.18 [0.14, 0.21] | 0.55 | 0.48 |
| Maximum neointimal thickness [mm] | 0.49 [0.30, 0.58] | 0.32 [0.23, 0.43] | 0.69 | 0.0055 |
| Lumen asymmetry | 0.06 [0.03, 0.08] | 0.04 [0.03, 0.06] | 0.63 | 0.062 |
| Fibrin ratio | 0.07 [0.05, 0.16] | 0.41 [0.31, 0.52] | 0.94 | $1.8 \times 10^{-10}$ |
| Fibrin score | 0.50 [0.29, 0.73] | 2.43 [1.94, 2.87] | 0.99 | $6.9 \times 10^{-13}$ |

**Figure 5.**
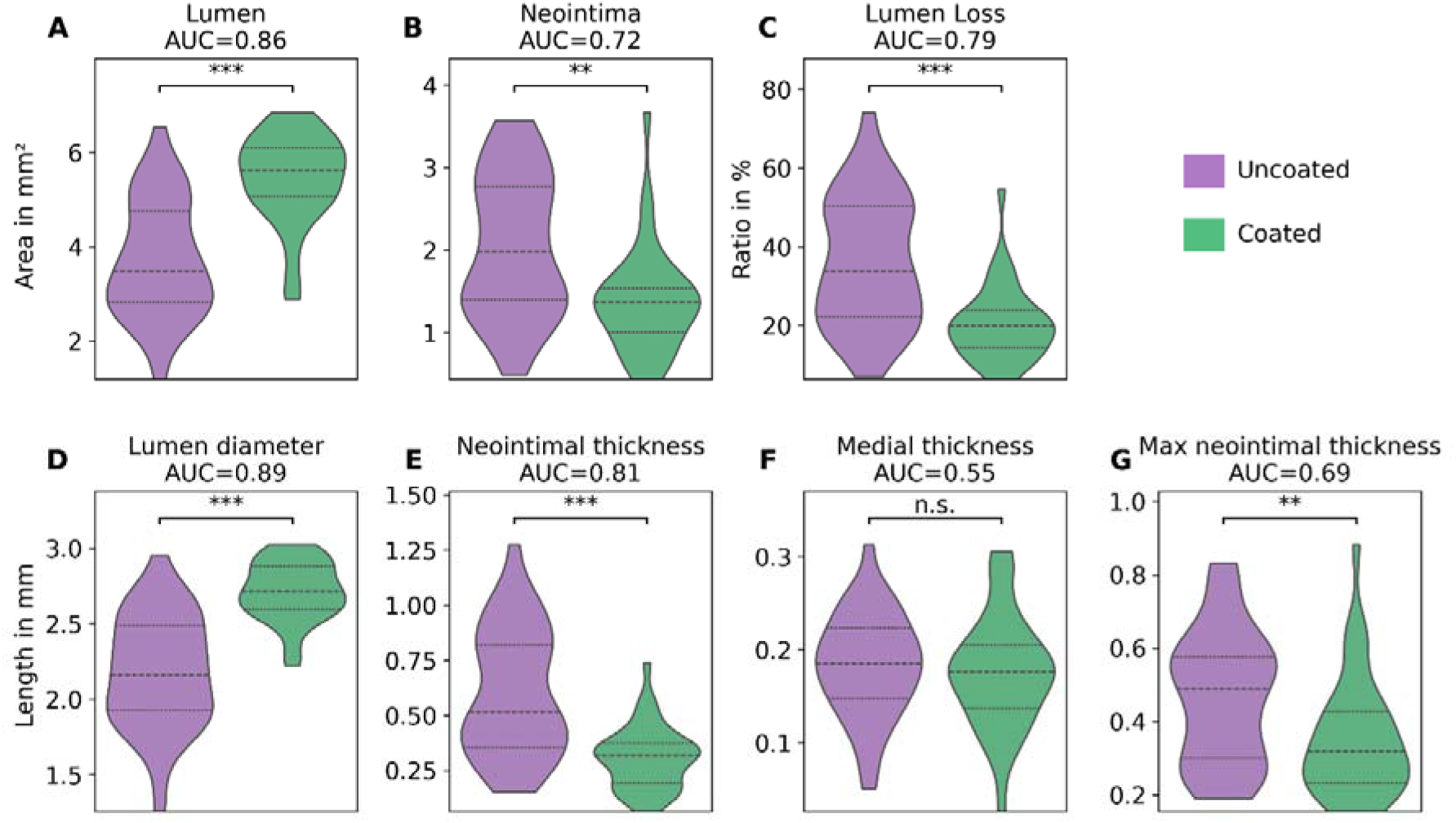
Classical morphology parameters derived by qHisto. Violin plots (36 coated/ 36 uncoated) for A) lumen area, B) neointima area, C) lumen loss, D) lumen diameter, E) neointimal thickness, F) medial thickness, and G) maximum neointimal thickness. *p < 0.05, **p < 0.01, ***p < 0.001. AUC: area under the curve; qHisto: quantitative histology.

In addition to classical metrics, improved parameters were assessed to further evaluate arterial remodeling in the cross-sections. These included the qHisto fibrin ratio and lumen asymmetry. The improved parameters were analyzed according to their ability in distinguishing between coated and uncoated balloons used for stenting (**Figure 6. Improved parameters derived by qHisto**. Violin plot (36 coated/ 36 uncoated) for A) lumen asymmetry, B) ratio of fibrin area to neointima area derived by qHisto, and C) fibrin score from conHisto. D) Correlation of fibrin ratio from qHisto vs. fibrin score from conHisto (R = 0.77, p = 3.61 × 10^-15^). *p < 0.05, **p < 0.01, ***p < 0.001.).

**Figure 6.**
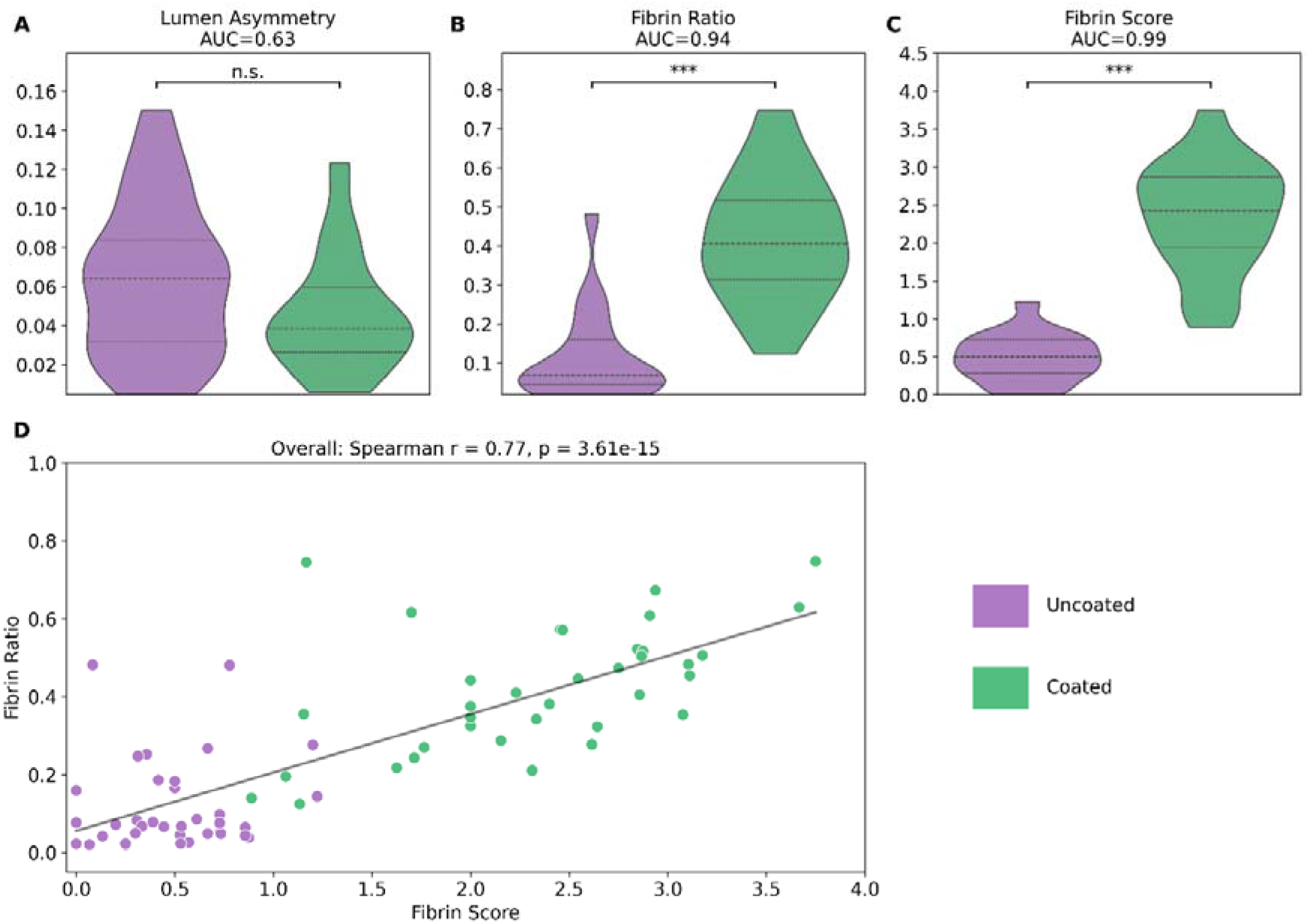
Improved parameters derived by qHisto. Violin plot (36 coated/ 36 uncoated) for A) lumen asymmetry, B) ratio of fibrin area to neointima area derived by qHisto, and C) fibrin score from conHisto. D) Correlation of fibrin ratio from qHisto vs. fibrin score from conHisto (R = 0.77, p = 3.61 × 10^-15^). *p < 0.05, **p < 0.01, ***p < 0.001. AUC: area under the curve; qHisto: quantitative histology; conHisto: conventional histology.

Both, the qHisto fibrin ratio and the conHisto fibrin score showed an excellent separation between coated and uncoated balloons (qHisto: AUC = 0.94, conHisto: AUC = 0.99, **Figure 6. Improved parameters derived by qHisto**. Violin plot (36 coated/ 36 uncoated) for A) lumen asymmetry, B) ratio of fibrin area to neointima area derived by qHisto, and C) fibrin score from conHisto. D) Correlation of fibrin ratio from qHisto vs. fibrin score from conHisto (R = 0.77, p = 3.61 × 10^-15^). *p < 0.05, **p < 0.01, ***p < 0.001. B and C). Moreover, both showed a strong and statistically significant correlation (R = 0.77, p < 0.0001, **Figure 6. Improved parameters derived by qHisto**. Violin plot (36 coated/ 36 uncoated) for A) lumen asymmetry, B) ratio of fibrin area to neointima area derived by qHisto, and C) fibrin score from conHisto. D) Correlation of fibrin ratio from qHisto vs. fibrin score from conHisto (R = 0.77, p = 3.61 × 10^-15^). *p < 0.05, **p < 0.01, ***p < 0.001. D). **Figure 7. Examples of fibrin detection using qHisto and conHisto**. Both slides show similar amounts of fibrin coverage, while conHisto showed very different values for both slides, qHisto assesses similar fibrin content. A) Both qHisto and conHisto assess the fibrin content similarly within their respective evaluation systems. B) A substantial amount of fibrin is located away from the strut voids in the neointima, which is not taken into account by conHisto, whereas qHisto includes fibrin coverage throughout the whole neointimal area, leading to very different assessments of fibrin coverage. shows two slides with similar fibrin coverage, along with the determined qHisto fibrin ratio and the conHisto fibrin score. The conHisto scores were highly divergent (2.87 vs 0.08), while the qHisto ratios showed similar values (0.50 vs 0.48).

**Figure 7.**
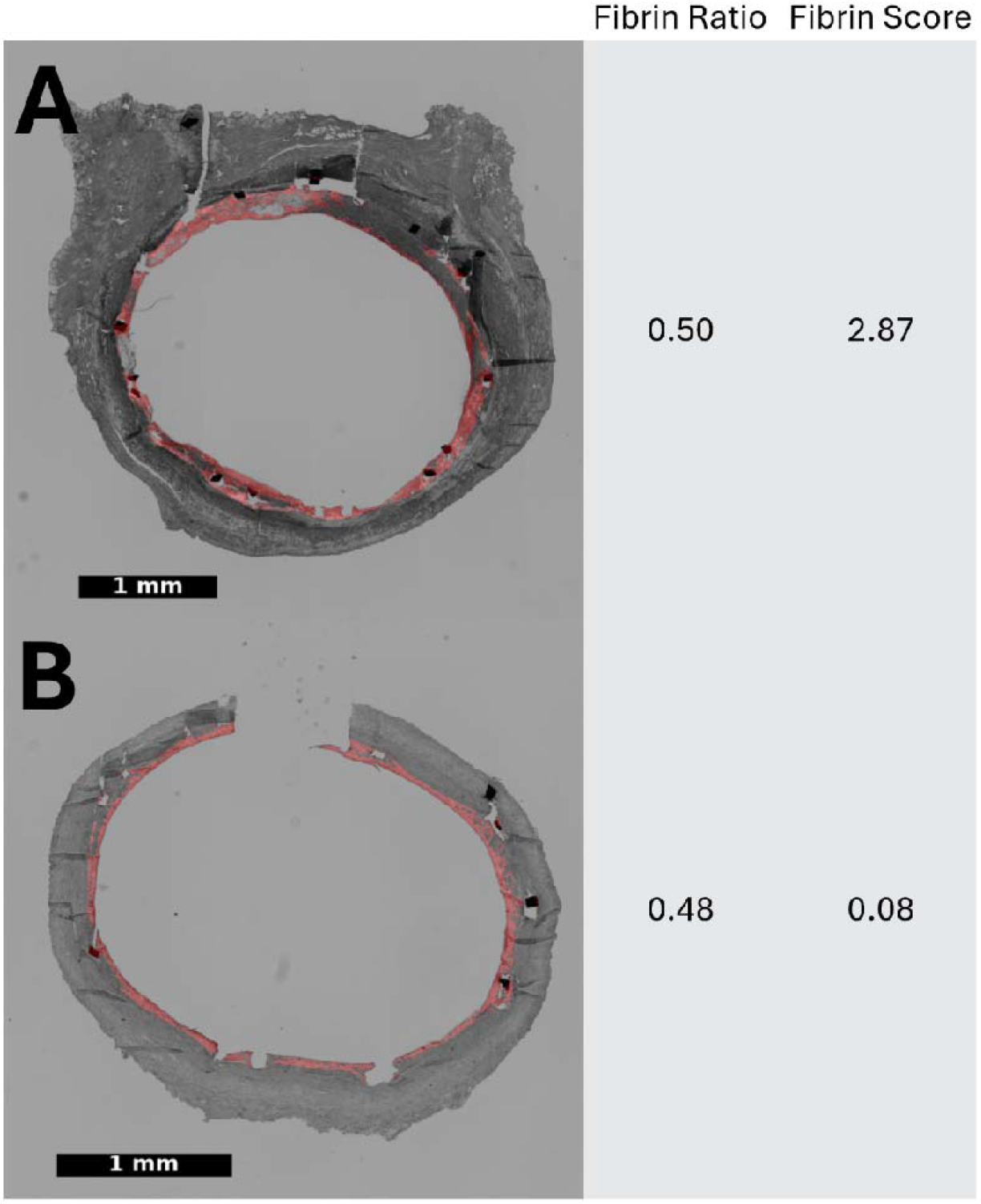
Examples of fibrin detection using qHisto and conHisto. Both slides show similar amounts of fibrin coverage, while conHisto showed very different values for both slides, qHisto assesses similar fibrin content. A) Both qHisto and conHisto assess the fibrin content similarly within their respective evaluation systems. B) A substantial amount of fibrin is located away from the strut voids in the neointima, which is not taken into account by conHisto, whereas qHisto includes fibrin coverage throughout the whole neointimal area, leading to very different assessments of fibrin coverage. qHisto: quantitative histology; conHisto: conventional histology.

The lumen asymmetry showed a moderate separation of coated and uncoated balloons (AUC = 0.63, **Figure 6. Improved parameters derived by qHisto**. Violin plot (36 coated/ 36 uncoated) for A) lumen asymmetry, B) ratio of fibrin area to neointima area derived by qHisto, and C) fibrin score from conHisto. D) Correlation of fibrin ratio from qHisto vs. fibrin score from conHisto (R = 0.77, p = 3.61 × 10^-15^). *p < 0.05, **p < 0.01, ***p < 0.001. A). **Figure 8** shows an example of low and higher lumen asymmetry (0.02 vs. 0.15) indicating strongly concentric and eccentric neointima formation, respectively.

**Figure 8.**
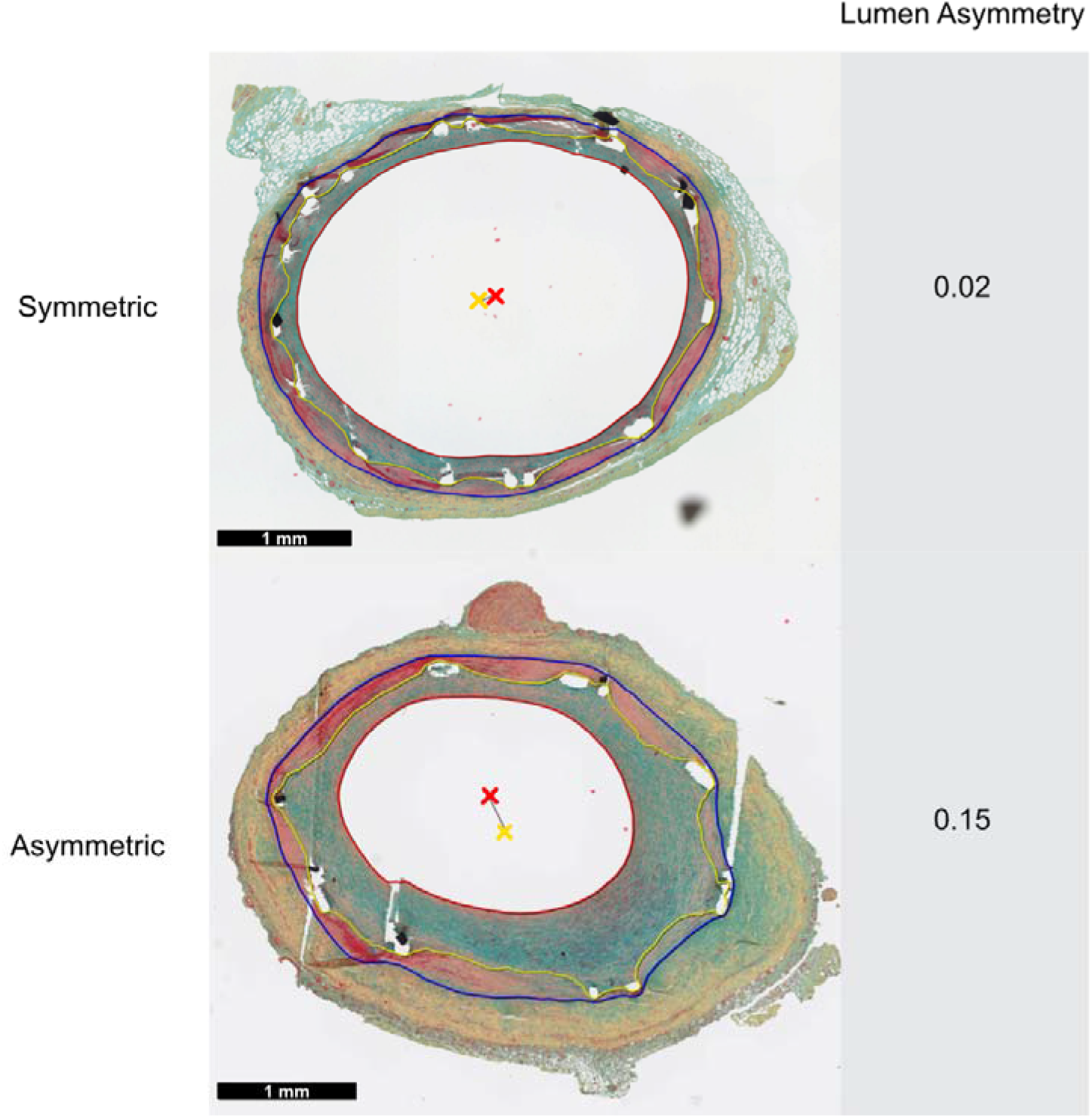
Examples for lumen asymmetry derived by qHisto. Lumen asymmetry shows low values for symmetric neointima formation (0.02) and higher values for asymmetric neointima formation (0.15). Crosses indicate the centroid of the lumen (red) and IEL (yellow). qHisto: quantitative histology; IEL: internal elastic lamina.

**Table 3** summarizes the results of the morphometric comparison of in-stent segments treated with uncoated and coated balloons. The medians of the lumen area, diameter, qHisto fibrin ratio, and conHisto fibrin score were lower in the samples treated with uncoated balloons. However, these samples showed higher median values for neointimal area, thickness, lumen loss, medial thickness, and maximum neointima than the samples treated with coated balloons. These results are consistent with expectations based on the known drug-related effects of treatment with coated balloons [16].

In addition to the diagnostic capability of qHisto, the time savings compared to manual morphometric analysis were analyzed. The entire pipeline took an average of 87 seconds per cross-section for the coating dataset. Compared to an average manual analysis time of 15 minutes, this corresponds to a time saving of 90%.

## 4. Discussion

In this work, we have successfully developed a fully automated system for the segmentation of arterial wall structures and strut voids, capable of achieving precise automated quantifications of morphometric parameters. Additionally, improved parameters were implemented to overcome limitations of previous parameters and thus improve the analysis of tissue alterations during different treatments. The system effectively reduces the time required for annotation and morphometric analysis by 90%, while providing objective and reproducible results. In comparison to another semi-automatic segmentation approach for arterial cross-sections, which achieved a time saving of around 80%, qHisto shows a higher reduction in analysis time [21]. Our pipeline is fully automated, enables a standardized, comprehensive analysis, and user interaction is limited to uploading data and reviewing the results.

To evaluate the predictions of the arterial structures the DSC was calculated and compared with the estimated variability between human operators. This demonstrates that the model achieves comparable median DSC values to the inter-operator variability (lumen: 0.996 vs. 0.997, neointima: 0.949 vs. 0.986, media: 0.892 vs. 0.905). The DSC values achieved by qHisto demonstrate a reliable segmentation of the respective structures, which is suitable for robust and accurate quantitative morphometric analysis. In comparison, models in other studies trained on H&E-stained scans of tissue-engineered vascular grafts achieved a lower DSC of 0.939 for the lumen and a marginally higher DSC of 0.907 for the media [26]. Furthermore, we have shown that the quality of the segmentation results is ideally suited for the morphological analysis of vascular remodeling processes induced by different treatment methods. The F1 score was calculated to evaluate the performance of the strut void segmentation model. A median F1 score of 0.97 was determined, demonstrating a strong balance between sensitivity and precision and confirming reliable detection of strut voids.

To evaluate the automated determination of the established morphometric parameters, the relative deviation from the manual assessment was calculated. The median difference in the calculated areas was close to 0%, while the structural diameters deviate within 4% for all structures. The maximum neointima thickness showed a median deviation of approximately 25%. However, manual determination of the length measurements, especially the maximum neointima thickness, is very prone to error, as a visual assessment is subjective to a high degree of bias and subjectivity. In contrast, qHisto enables an objective and reproduceable assessment through mathematically precise automatic calculation.

The implementation of an automatic determination of the fibrin area within the neointima enables a quantitative fibrin detection, thus overcoming limitations of the previous semi-quantitative conHisto fibrin score which considers only the fibrin area surrounding the strut voids as demonstrated in **Figure 7. Examples of fibrin detection using qHisto and conHisto**. Both slides show similar amounts of fibrin coverage, while conHisto showed very different values for both slides, qHisto assesses similar fibrin content. A) Both qHisto and conHisto assess the fibrin content similarly within their respective evaluation systems. B) A substantial amount of fibrin is located away from the strut voids in the neointima, which is not taken into account by conHisto, whereas qHisto includes fibrin coverage throughout the whole neointimal area, leading to very different assessments of fibrin coverage. [8]. Considering fibrin independently of the stent struts not only proposes a standardized, objective method, but also makes fibrin detection possible in non-stented sections. However, in some slides of lower quality, this method tends to overestimate the fibrin content. This is particularly noticeable in regions with dark contrast, such as folds and incompletely dislodged stent struts that have become lodged outside the strut voids. Nevertheless, the full-automatic qHisto fibrin ratio correlates strongly positively with the manual conHisto score (R = 0.77, **Figure 6. Improved parameters derived by qHisto**. Violin plot (36 coated/ 36 uncoated) for A) lumen asymmetry, B) ratio of fibrin area to neointima area derived by qHisto, and C) fibrin score from conHisto. D) Correlation of fibrin ratio from qHisto vs. fibrin score from conHisto (R = 0.77, p = 3.61 × 10^-15^). *p < 0.05, **p < 0.01, ***p < 0.001. D). Furthermore, both the qHisto fibrin ratio (AUC = 0.94, **Figure 6. Improved parameters derived by qHisto**. Violin plot (36 coated/ 36 uncoated) for A) lumen asymmetry, B) ratio of fibrin area to neointima area derived by qHisto, and C) fibrin score from conHisto. D) Correlation of fibrin ratio from qHisto vs. fibrin score from conHisto (R = 0.77, p = 3.61 × 10^-15^). *p < 0.05, **p < 0.01, ***p < 0.001. B) and the conHisto fibrin score (AUC = 0.99, **Figure 6. Improved parameters derived by qHisto**. Violin plot (36 coated/ 36 uncoated) for A) lumen asymmetry, B) ratio of fibrin area to neointima area derived by qHisto, and C) fibrin score from conHisto. D) Correlation of fibrin ratio from qHisto vs. fibrin score from conHisto (R = 0.77, p = 3.61 × 10^-15^). *p < 0.05, **p < 0.01, ***p < 0.001. C) show a strong separation between coated and uncoated balloons. Therefore, the qHisto fibrin ratio is suitable for assessing vascular healing processes after stent insertion with different coated balloons [31].

The automated determination of the lumen asymmetry enables the assessment of neointima concentricity in in-stent restenosis. It overcomes the limitation of the previously proposed ratio of minimum to maximum neointima thickness [28], which, in case of thin neointima, can lead to a misinterpretation of the medical relevance of high asymmetry. In contrast, lumen asymmetry is based solely on the centroids of the lumen and the IEL and therefore represents a stable morphometric parameter for assessing the symmetry of in-stent neointimal growth [22].

The qHisto artery pipeline has been integrated into the BIOQIC Apps for internal use at Charité. This makes it easily accessible for daily preclinical research and enables efficient analysis of large datasets through parallel processing on the Charité/BIH HPC cluster.

We have demonstrated that qHisto is highly suitable for the automated segmentation and morphometric quantification of porcine stented arterial cross-sections. The automatic and time-efficient pipeline opens up several perspectives for future investigations. For example, the qHisto artery pipeline could be integrated into routine preclinical workflows. Future studies could explore how this method can be applied to other datasets, such as non-stented sections or sections from other organisms like mice, rats, or rabbits. Moreover, parameters such as lumen asymmetry could be investigated in greater detail regarding their diagnostic relevance by analyzing cohorts based on the extent of symmetrical or asymmetrical neointima formation.

## Limitations

The segmentation results generally showed very high accuracy. In exceptional cases, however, manual correction is required to improve precision. In our test dataset, this only affected one image, which corresponds to less than one percent of the processed images. Furthermore, this image represents a highly complex scenario that can be challenging even for manual assessment. Nevertheless, a manual review of the results is essential to ensure high quality and minimize errors.

## Conclusion

In this paper, we presented a deep learning framework for automated segmentation and morphometric analysis of arterial cross-sections. The system, which extends the qHisto framework to arterial cross-sections, determines established morphological key parameters in a standardized and reproduceable way while also introducing improved parameters. The qHisto artery pipeline provides segmentation results of comparable quality to manual segmentation while reducing analysis time by 90%.

Furthermore, qHisto reliably distinguished between arterial segments treated with coated and uncoated balloons, thus demonstrating its ability to detect biologically relevant treatment effects. Consequently, the qHisto pipeline for arterial cross-sections enables a comprehensive, standardized and time-efficient assessment of morphological parameters in preclinical research settings.

## Acknowledgements

We thank the German Research Foundation (DFG) for funding (KS, HT, TM to CRC1340 Matrix-in-Vision; KS and TM to FOR5628; TM, RB and NJ to BIOQIC GRK 2260 and Sa901/33-1 M5). Open access funding enabled and organized by Projekt DEAL.

## Data availability

The corresponding author will make the data available upon reasonable request. The source code underlying this study will be made publicly available upon publication under the following repository: https://gitlab.com/Charite-IMI/qhisto_arteries.

